# The Origin of Coupled Chloride and Proton Transport in a Cl^−^/H^+^ Antiporter

**DOI:** 10.1101/064527

**Authors:** Sangyun Lee, Heather B. Mayes, Jessica M. J. Swanson, Gregory A. Voth

## Abstract

The ClC family of transmembrane proteins functions throughout nature to control the transport of Cl^−^ ions across biological membranes. ClC-ec1 from *Escherichia coli* is an antiporter, coupling the transport of Cl^−^ and H^+^ ions in opposite directions and driven by the concentration gradients of the ions. Despite keen interest in this protein, the molecular mechanism of the Cl^−^/H^+^ coupling has not been fully elucidated. Here, we have used multiscale simulation to help identify the essential mechanism of the Cl^−^/H^+^ coupling. We find that the highest barrier for proton transport (PT) from the intra- to extracellular solution is attributable to a chemical reaction—the deprotonation of glutamic acid 148 (E148). This barrier is significantly reduced by the binding of Cl^−^ in the “central” site (Cl^−^_cen_), which displaces E148 and thereby facilitates its deprotonation. Conversely, in the absence of Cl^−^_cen_ E148 favors the “down” conformation, which results in a much higher cumulative rotation and deprotonation barrier that effectively blocks PT to the extracellular solution. Thus, the rotation of E148 plays a critical role in defining the Cl^−^/H^+^ coupling. As a control, we have also simulated PT in the ClC-ec1 E148A mutant to further understand the role of this residue. Replacement with a non-protonatable residue greatly increases the free energy barrier for PT from E203 to the extracellular solution, explaining the experimental result that PT in E148A is blocked whether or not Cl^−^_cen_ is present. The results presented here suggest both how a chemical reaction can control the rate of PT and also how it can provide a mechanism for a coupling of the two ion transport processes.

## Introduction

The ClC channels and transporters constitute a large and intriguing family of transmembrane proteins, including both chloride channels and chloride/proton antiporters.^1^ They are found in a wide range of organisms, including many prokaryotes and nearly all eukaryotic cells.^2-4^ Different isoforms are involved in many different physiological functions, such as stabilization of the membrane potential (ClC-1), regulation of transepithelial Cl^−^ transport (ClC-2, -Ka, and -Kb), ion homeostasis of endosomes (ClC-3,4,5, and 6), lysosome acidification (ClC-7), and acid resistance in bacterial cells (ClC-ec1).^2,5,6^ Defects in ClC proteins are known to cause several hereditary diseases, such as myotonia congenita, Dent’s disease, Bartter’s syndrome, osteopetrosis, and idiopathic epilepsy.^1,3,6^

ClC-ec1, a bacterial ClC transporter from *Escherichia coli*, mediates the exchange (antiporting) mechanism of Cl^−^ and H^+^ ions through the membrane (Figure 1A). It utilizes a secondary active transport mechanism in which a concentration gradient of either Cl^−^ or H^+^ drives the transport of the other ion, as confirmed by multiple studies employing a wide range of concentration gradients for Cl^−^ and H^+^.^5,7^ Transport can occur in either direction, with one of the two directions shown in Figure 1B. The Cl^−^:H^+^ exchange ratio (~2:1) is consistent within a wide range of concentration gradients of both ions, suggesting that the Cl^−^ and the H^+^ fluxes in the ClC-ec1 are strongly coupled.^7,8^ Later experiments ^9,10^ directly measured the turnover rate of the Cl^−^ efflux out of the liposome while there is the H^+^ influx against the pH gradient, and confirmed that the Cl^−^/H^+^ exchange ratio of the ClC-ec1 is (2.2 ± 0.1):1.^11^

**Figure 1.**
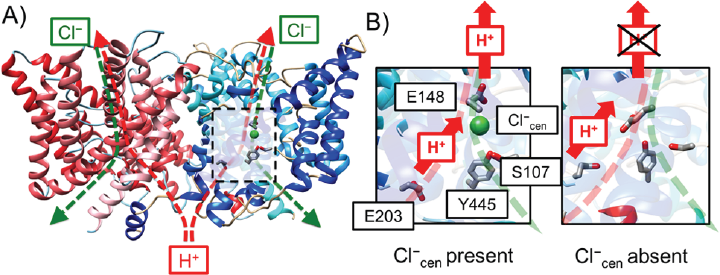
(A) Overview of the structure of the ClC-ec1 antiporter and transport pathways for Cl^−^ (green dashed) and H^+^ (red dashed) based on PDB ID: 1OTS.^12^ ClC-ec1 is a homodimer (monomer A shown in blue and monomer B in red). The central region of monomer A is highlighted by the dashed black box. (B) Schematic picture of the PT pathway with S_cen_ either occupied (Cl^−^_cen_ present, left) or unoccupied (Cl^−^_cen_ absent, right) by a chloride ion. The H^+^ flux is represented as red arrows, with positive flux defined as transport from the intracellular to the extracellular solution. The “X” over the upper H^+^ on the right indicates that no PT to the extracellular bulk is observed when Cl^−^_cen_ is absent.

To reveal important residues for proton transport (PT), site-directed mutagenesis experiments have targeted several Glu and Asp residues^13,14^. These studies showed that H^+^ flux was blocked while Cl^−^ flux was still observed in the E148A and E203Q mutants. In addition, the Cl^−^ uptake rate was increased at low pH in the E203Q mutant, similar to WT, but became pH-independent in the E148A mutant. Interestingly, while no proton flux was observed for E148A with or without Cl^−^ in the system, Feng *et al.*^15^ found that adding free glutamate to the solution rescued proton flux across the membrane, providing additional clues as to the possible PT transfer mechanism. Several key steps in the Cl^−^/H^+^ exchange process were proposed based on these and other experimental findings: 1) E148 (Glu_ex_) and E203 (Glu_in_) participate in the PT process, 2) protonation of E148 opens the extracellular gate and allows Cl^−^ transport, and 3) the Cl^−^ and H^+^ transport pathways overlap from E148 to the extracellular solution, as shown in Figure 1, but diverge below E148.^12,13^ As previously noted, transport can occur in the direction shown in Figure 1B or the opposite direction, and researchers have proposed fully reversible transport mechanisms.^11,16,17^

While these studies and others provided crucial insight into the exchange mechanism, remaining uncertainties resulted in different proposals for the elementary steps.^1^ For example, some researchers proposed that PT in the central region occurs with Cl^−^ occupying the central site (S_cen_),^17^ while others proposed PT occurs without Cl^−^ at S_cen_ (Cl^−^_cen_).^11,16,17^ This question prompted our previous study of PT in the central region, in which we found that while the barrier for PT from E203 to E148 was lower with Cl^−^ present, but the calculated rate constants for both cases were significantly faster than the measured turnover rate.^18^ This study raised the obvious question of which PT step would be rate-determining and how could PT be coupled to Cl^−^ transport, thus motivating the present study.

The full PT pathway through the protein includes transit beyond the central region: 1) from the solution on the intracellular side of the protein to E203, and 2) from E148 to the solution on the extracellular side. The latter step is more likely to be coupled with Cl^−^ since the Cl^−^ and H^+^ transport pathways fully overlap in this region (Figure 1B), while E203 is separated from the central Cl^−^ binding site by ~ 10 Å. Moreover, unlike E148, residue E203 is not strictly conserved in CLC, suggesting that its function is less critical.^13-15^ Thus, herein we focus on PT from E148 to the extracellular solution and assume that the rate of step 1 is relatively fast. Using enhanced free energy sampling coupled with multiscale reactive molecular dynamics (MS-RMD),^19-22^ we calculate the free energy profiles (potentials of mean force, PMFs) for PT from E148 to the extracellular bulk in the presence and absence of the Cl^−^_cen_. We show that this step has a rate constant that is similar to that inferred from the overall measured PT rate, suggesting that it is rate-determining during PT from the intra- to extracellular bulk. However, the barriers are asymmetric with respect to directionality, and the smallest calculated rate constant for PT from the extra- to intracellular bulk is for transport from E148 to E203. Thus, in either direction, E148 deprotonation is likely rate limiting for PT and, as we will show later in this paper, this step is significantly facilitated by the presence of Cl^−^_cen_. We further identify an essential mechanism of Cl^−^/H^+^ coupling: in the absence of Cl^−^_cen_, E148 is stabilized in the down conformation, effectively blocking PT from intra- to extracellular solution, thus confirming a hypothesis put forth by Feng et al.^15^

As mentioned above, experiments^7,13,17^ have shown that the E148A mutant cannot transport protons, but it allows pH-independent Cl^−^ flux. To help explain this puzzling result, PMFs for PT from E203 to extracellular bulk were also calculated in the E148A mutant both in the presence and absence of Cl^−^_cen_. Our results show that PT past A148 is effectively blocked for both cases, in agreement with experimental findings. Since the residues near E148 are mainly hydrophobic the extracellular water molecules are separated from those that can fill the central region in the WT system. E148 transfers a proton through this region by rotating its side chain from the central waters to the external waters. However, the cavity near A148 in the E148A mutant remains dehydrated and the barrier for the hydrated excess proton to pass by the unprotonatable alanine residue is greatly increased, becoming effectively insurmountable over any physiologically relevant pH range.

## Methods

The details for the system setup and the parameterization of MS-RMD model are described in detail in the Supporting Information (SI). Briefly, the system is based on the ClC-ec1 dimer structure (PDB ID: 1OTS)^12^ and modeled with the CHARMM forcefield.^23,24^ The simulation was performed with the RAPTOR software^19^ to implement the MS-RMD description of PT, interfaced with the LAMMPS MD package (http://lammps.sandia.gov).^25^ Initial configurations for the simulations were obtained from a previous study of this system.^18^

### E148 rotation and deprotonation reaction paths

The PT mechanism from E148 to the extracellular solution was studied by calculating the PMFs for a two-step process: the rotation of the E148 side chain from its down to up conformation, followed by the deprotonation of E148 to the extracellular solution through intervening water molecules. The two steps were described by a single continuous collective variable (CV), which was the curvilinear pathway of the protonic center of the excess charge (CEC), following similar procedures previously described^22^ with additional details provided in the SI. Briefly, the path was identified by adding biases along the z-axis according to the metadynamics algorithm^26,27^ implemented in the PLUMED package,^28^ with wall potentials preventing sampling regions far from the protein pores. The curvilinear pathways with either Cl^−^_cen_ present or absent are shown in Figure 2. We note that the channel pore size is narrow at E148, but gradually increases as it goes to the extracellular solution. At the region above E148, the helical kink in panel B indicates a more complex pathway, where the excess proton migrates through various water molecules to the extracellular solution. However, when converging the PMF in the subsequent umbrella sampling (described next), a cylindrical confining potential was applied to confine the sampling space to the most relevant region as the pore size increases.

**Figure 2.**
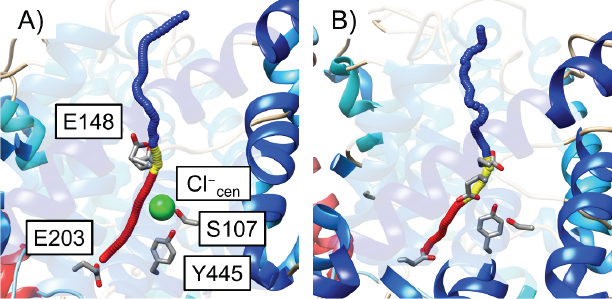
The curvilinear PT pathway for the CEC when Cl^−^_cen_ is present (A) or absent (B): PT from E203 to E148 (red), the rotation of protonated E148 (yellow), and PT from E148 to the extracellular side (blue). E148 is shown in the “up” conformation on the left (A) and, in the “down” conformation on the right (B).

### E148 rotation and deprotonation PMF calculations

The conformations along each PT pathway were sampled using the replica exchange umbrella sampling (REUS) method.^29^ Windows were separated by 0.25 Å in the z direction of the CV, defined as the distance of the CEC from E148 along the curvilinear pathway described in the previous subsection, with the direction of the harmonic umbrella potential defined by the tangent vector of the path at the window center.^27^ The force constant of the harmonic potential was set to be 30 *kcal*·mol^−^·Å^−2^. A cylindrical wall potential with a 5 Å radius was added to the direction perpendicular to the pathway as the proton entered bulk solution in line with previous ion channel PMF studies.^30,31^ The PMFs were calculated using the WHAM algorithm^32^ combining regions the MS-RMD models, as described in the SI. Additional features of this PMF calculations are also described in the SI.

### E148A reaction path and proton transport PMF calculations

The PMF for PT in E148A mutant was calculated with a similar procedure as that used for WT, but with a wider range for the CV: where the excess proton is transferred from E203, through the central region via water molecules, past A148, and then to the extracellular solution. We employed the MS-RMD model for E203 from our previous work^18^ and A148 was treated by the CHARMM classical force field. The initial configurations for the metadynamics simulations to obtain the curvilinear paths were obtained from the WT simulations, after mutating residue 148 to alanine and equilibrating with classical MD for 1 ns. Then the PMF for PT from E203 to the extracellular solution was calculated from REUS along the curvilinear path determined from the metadynamics simulations, with both Cl^−^_cen_ present and absent, consistent with the procedure described for the WT protein.

### Proton transport rate constants and pK_a_ calculations

The PT rate constants were estimated using transition state theory as follows,^18,33^

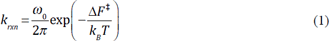

where *k_B_* is Boltzmann’s constant, *T* is the simulation temperature (300 K), and Δ*F*^‡^ is the free energy barrier height in the PMF. The fundamental frequency *ω*_0_ is that of the reactant state oscillations around its minimum, which is defined as

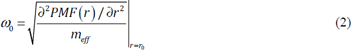

where *r*_0_ is the local minimum in the PMFs. The effective mass of the excess proton CEC, *m_eff_*, was determined using the equipartition theorem, 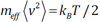, where the value of 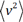 was calculated from the MS-RMD trajectory sampled at *r*_0_.

The pK_a_ of E148 was estimated using the equation for calculating the equilibrium constant of binding of the substrate at the binding site of the protein, based on the one-dimensional PMF for the substrate moving along the channel axis with the cylindrical potential applied at the channel entrance:^34^

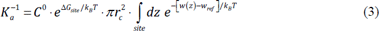

where the substrate is the excess proton and the binding site is E148. Here, *C°* is the standard state concentration (1 M = 1/1660 Å^−3^), and Δ*G_site_* is the free energy cost introduced by the cylindrical potential at the substrate binding site (the CEC is at E148.). The value of Δ*G_site_* is zero in this case, because the sampling area for the CEC at E148 is smaller than the radius of the cylindrical potential, and no bias is felt by the CEC at this region. The quantity *r* is the radius of the cylindrical potential, which is set to be 5 Å. The quantity *w(z)* is the one-dimensional PMF as a function of the CV, *z*, which is the distance of the CEC along the curvilinear pathway, while *w_ref_* is the asymptotic value of *w*(*z*), when the excess proton is at a long distance away in the extracellular solution. When the Boltzmann factor of *w(z)* is integrated, the lower boundary for *z* is placed at the position of E148. The pKa of E148 is insensitive to the choice of the upper boundary for *z*, since the Boltzmann factor of *w*(*z*) is quickly converged as the *z* value goes to the extracellular solution; the pKa of E148 changes only 0.001, when the upper boundary is set to be any *z* value between 2 Å above the lower boundary and the extracellular solution.

## Results and Discussion

### Proton transport between the central region and extracellular bulk water region

The PMFs for PT from E148 in the central region to the extracellular solution with Cl^−^_cen_ either present or absent (Figure 3) reveal that PT in this region occurs via a two-step process: 1) the change of the orientation of E148 side chain from the down to the up conformation, and 2) the deprotonation of E148 in the up conformation followed by PT to the extracellular solution. The structures of the down and up minima are shown in Figure 4. In the down orientation (Figure 4A and C), the carboxyl group of E148 is hydrogen bonded to water molecules in the central region, with either Cl^−^_cen_ present or absent. To move to the up conformation (Figure 4B and D), the carboxyl group breaks the hydrogen bonds with the water molecules in the pore (corresponding with the barrier in the PMF between the two local minima) and then makes new hydrogen bonds with the water molecules from the extracellular solution. Thus, E148 both separates the water molecules in the pore from those leading to extracellular solution, and acts as a bridge for the excess proton to cross this region. Following the rotation of the protonated E148 side chain, E148 must deprotonate (surmounting an additional energy barrier) to complete the transfer to the extracellular bulk water.

**Figure 3.**
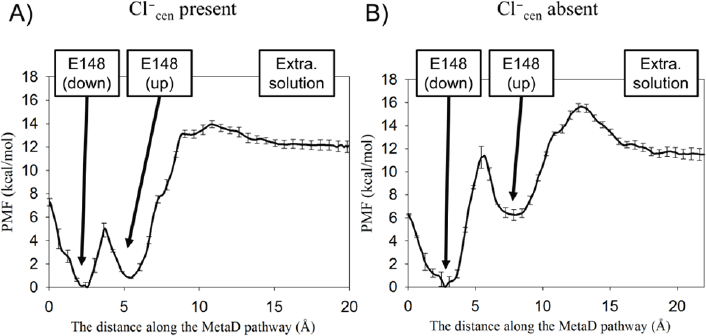
The PMF for a two-step PT process with Cl^−^_cen_ present (A) or absent (B), including the rotation of E148 from the down to the up conformations, followed by the deprotonation of E148 to the extracellular solution.

**Figure 4.**
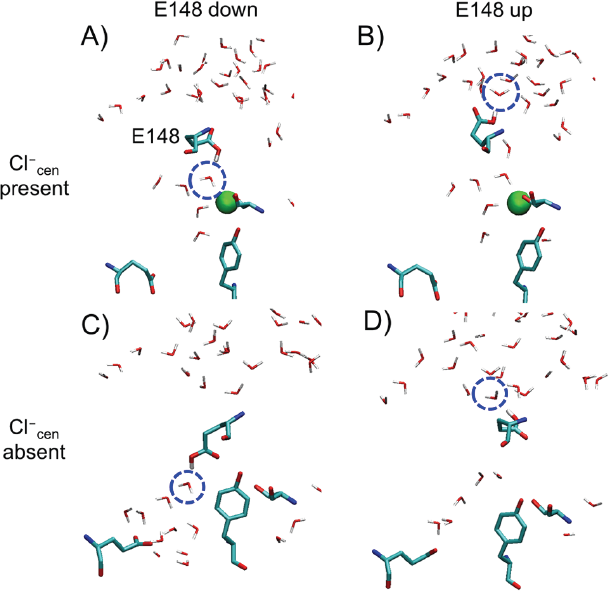
Representative configurations for the local energy minima of the PMFs in Fig. 2, with Cl^−^_cen_ present (A, B) or absent (C, D), and with E148 in the down (A, C) or the up conformation (B, D).

Both the energy well depth and position of the protonated side chain in the pore differ between the Cl^−^_cen_ present and absent cases. As shown in Figure 4C, when Cl^−^_cen_ is absent and E148 is down it occupies the vacated central site (S_cen_) and is 6.3 kcal/mol more favored than the up conformation. When Cl^−^_cen_ is present, it sterically prevents E148 from occupying S_cen_, keeping the E148 up/down conformational and energy change relatively small. We also calculated the PMF for rotation of deprotonated E148 in the absence of Cl^−^_cen_ (Figure S3), which showed that the down conformation of E148 (where the negatively charged side chain gets close to S_cen_) is ~10 kcal/mol energetically more favorable than the up conformation. When E148 is protonated, the down conformation in the absence of Cl^−^_cen_ is stabilized by only ~ 6 kcal/mol. The greater stabilization of the negatively charged state of E148 is consistent with a previous computational study^35^ that calculated the electrostatic potential energy profile along the Cl^−^ pathway, finding that Cl^−^ at the S_cen_ site is stabilized by a surrounding net positive charge.

X-ray crystal structures can represent snapshots of a protein’s conformational change at different intermediate states. Thus, Figure S4 compares the simulation intermediates found herein to three different crystal structures: WT of ClC-ec1 (PDB ID: 1OTS)^12^, E148Q mutant of ClC-ec1 (1OTU),^12^ and WT of cmClC (3ORG).^16^ These crystal structures capture different conformations of E148 and different anion occupancy in the external, central, and internal sites (S_ext_, S_cen_, and S_int_). Residue Q148 in the E148Q mutant is considered a mimic of the protonated state of E148 in WT. The 1OTU crystal structure (Cl^−^_cen_ present) overlaps well with the simulation structure taken from the window at the local energy minima for up conformation of E148 in the PMF with Cl^−^_cen_ present. The WT crystal structure 1OTS (Cl^−^_cen_ present) overlaps well with down confirmation from the same PMF. Since the two conformations are nearly isoenergetic, it is not surprising that E148Q aligns better with the E148-up simulation conformation. Finally, 3ORG (Cl^−^_cen_ absent) overlaps well with the simulation structure of E148 in the down conformation from the PMF with Cl^−^_cen_ absent. The E148 up conformation with Cl^−^ _cen_ absent is a higher energy state that is unlikely to be captured in a crystal structure.

The presence of Cl^−^_cen_ changes not only the dominant conformations of E148, but also the energetics of rotation and deprotonation. Focusing first on the deprotonation of E148 toward extracellular solution, the PMFs plateau at *x* > 16 Å along the pathway (CV), where the excess proton is no longer interacting with the protein. The height of the free energy barrier for the second step (deprotonation) is higher with Cl^−^_cen_ present (13.1 kcal/mol), compared to that with Cl^−^_cen_ absent (9.3 kcal/mol). Since S_cen_ site is ~ 4 Å below E148, it follows that deprotonation (excess proton moving away from Cl^−^_cen_) will be more difficult in the presence of Cl^−^_cen_. Note that in the opposite direction (extra- to intracellular) the opposite is true, as shown in our previous work.^18^ Since the excess proton moves toward S_cen_ during PT from E148 to E203, the presence of Cl^−^_cen_ facilitates E148 deprotonation. Once the cost of rotation is factored in, the presence of Cl^−^_cen_ also facilitates PT from E148 to extracellular solution. The total free energy difference between the minimum in the PMFs in Figure 3 (protonated E148 in the down position) and the maximum (deprotonation of E148 in the up position), is higher with Cl^−^_cen_ absent (15.7 kcal/mol) than with Cl^−^_cen_ present (13.5 kcal/mol). The reason the presence of an anion in one position (at S_cen_) can have the same facilitating effect on PT in opposite directions is due to the rotation of E148 and steric competition between Cl^−^ and E148 for S_cen._ As discussed earlier, the down conformation of E148 is energetically favored in the absence of Cl cen, but the down rotation of E148 is sterically blocked by the presence of Cl^-^_cen_, minimizing the cost of E148 rotation from the pore-facing (‘down’) to the extracellular-facing (‘up’) conformation.

The effective rate constant, *k_eff_,* was calculated to obtain the rate constant of the two-step (rotation and deprotonation) process: *k_eff_ =k_2_·k_1_/k*_−1_, assuming that the first step reaches the equilibrium compared to the second *(k_2_ = k*_−1_), where *k_1_* and *k_−1_* are the forward and the backward rate constants for the first step in the PMF, and *k_2_* is for the forward rate constant for the second step. Table 1 shows that *k_eff_* with Cl^−^_cen_ present is 0.81 *ms*^−1^, and with Cl^−^_cen_ absent is 7.7×10~^3^ *ms^−1^. k_eff_* with Cl^−^_cen_ present is comparable to the experimental value of the turnover rate for the overall PT process, 1.0 *ms*^−1 11,14^, (calculated using the Cl– turnover rate of 2.3 *ms*^−1^ and the Cl^−^:H^+^ exchange ratio of 2.2:1). Thus, when the overall PT process is described in the direction from the intra- to extracellular side of the protein, as shown in Figure 1B, PT from E148 to the extracellular region with Cl^-^_cen_ present is a likely candidate for the rate-limiting step for the overall PT process. In contrast, *k_eff_* with Cl^−^_cen_ absent is on the order of sec^−1^, which may be too slow to be measured in conventional experimental techniques, as it would be difficult to separate from the background leak current through the membrane.^8,9,14,36^

**Figure 5.**
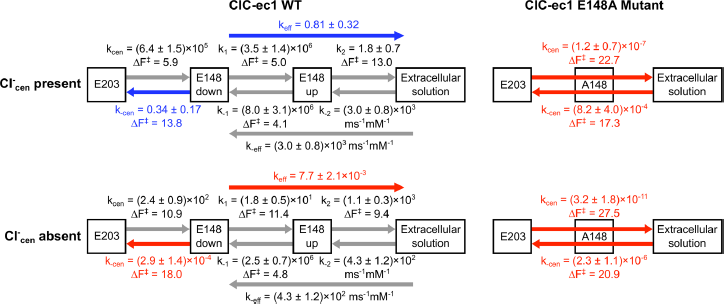
Schematic representation of the PT mechanism in ClC-ec1 WT (left) and its E148A mutant (right), with Cl^−^_cen_ present (top) or absent (bottom). The arrows show PT steps, with k_eff_ calculated as described in the text. The arrow direction indicates the direction of the H flux. The free energy barriers (in *kcal/mol)* and corresponding rate constants (in *ms*^−1^ unless otherwise indicated) are shown above or below the arrows (for outward and inward flux, respectively). Steps originating from the extracellular solution have second order rate constants. The gray arrows represent fast (non-limiting) steps, blue represent rate-limiting steps, and red represent steps that effectively block PT. The rate constants for H^+^ flux between E203 and E148 down were calculated from PMFs in a previous study. Errors in the rate constants were estimated by calculating it in four consecutive blocks in the trajectories for each window. The experimental value for the turnover rate for PT is 1.0 *ms*^−1^·^10,11^

It is known that the Cl^−^/H^+^ exchange mechanism can operate in both directions,^13,37^ where the overall H^+^ flux goes from the intracellular to the extracellular side of the protein (outward H^+^ flux) or in the opposite direction (inward H^+^ flux), depending on the directionalities of the concentration gradients of Cl^−^ and H^+^. The energy barriers for PT between the central region and the extracellular solution (Figure 3) and between E148 and E203 in the central region (our previous study) are highly asymmetric. For the outward H^+^ flux (the direction shown in Figure 1B), our previous study^18^ showed that PT from E203 to E148 is unlikely to be rate-limiting, regardless of the presence of Cl^−^_cen_. This study indicates that the combined PT steps of E148 rotation and deprotonation to the extracellular solution are likely rate-limiting for outward proton flux, facilitated by Cl^−^_cen_. Figure 5 shows a schematic representation of the PT mechanisms in both directions with the calculated rate constants each step.

However, the rate-limiting steps are likely reversed in the opposite direction. For the inward H^+^ flux, the PT rate constant from E148 to E203 with Cl^−^_cen_ present is 0.34 *ms*^−1^ (Figure 5), which is comparable to the experimental PT turnover rate. With Cl^−^_cen_ absent, it is 2.9×10~^4^ *ms*^−1^ decreasing PT from E148 to E203 below detectable levels. For PT from the extracellular solution to the central region (right to left in Figure 3), the *k_eff_* is estimated at 3.0×10^3^ *ms*^−1^ · *mM*^−1^ with Cl^−^_cen_ present and 4.3×10^2^ *ms*^−1^·*mM*^−1^ with it absent, which are second order rate constants depending both on protein binding site availability and the proton concentration in the extracellular solution. Thus, for the inward H flux, PT from E148 to E203 has the smallest rate constant and is again facilitated by Cl^−^_cen_.

The pKa of E148 was calculated using Eq. 3 at the local energy minima in the PMF for the up and the down conformations of E148. The pKa of E148 with Cl _cen_ present is 6.9 when E148 is in the down conformation, and 6.4 for the E148 up conformation. The pKa of E148 with Cl^−^_cen_ absent is 6.8 for down and 2.6 for the up conformation. As previously noted, the E148 up Cl^−^_cen_ conformation with Cl^−^_cen_ absent represents a high energy state that does not significantly contribute to the ensemble of states and thus contributes little to the overall proton binding affinity of E148. The pKa values at other conformational states are comparable to the experimental pKa value of 6.2,^38^ providing validation of the PMF s presented here.

### Proton transport in the E148A mutant

PT was also simulated for the ClC-ec1 E148A mutant with Cl^−^_cen_ both present and absent, where the excess proton is transferred from E203, through the central region, and to the extracellular solution. The PMFs for E148A mutant show that the free energy barrier is decreased with Cl^−^_cen_. present by 5.1 kcal/mol (Figure 6). This difference is similar to that in the PMFs for WT in the central region, where the free energy barrier for PT from E203 to E148 is decreased by 5.0 kcal/mol.^18^

In the WT protein, the excess proton is transferred through the narrow region above S_cen_ by E148 — while protonated E148 rotates between the central region and the extracellular solution. However, in the E148A mutant, A148 is non-protonatable and the region around A148 is narrow and dehydrated. Therefore, the free energy cost required for the excess proton transfer to the extracellular solution is greatly increased. The free energy maxima in the PMFs correspond to the point at which the excess proton is located in a narrow pore near A148. The PMFs with Cl^−^_cen_ present or absent show that the free energy barriers are high enough to reduce PT to lower than background levels in both outward and inward H^+^ fluxes, and regardless of the presence of Cl^−^_cen_. Our results agree with the experimental finding^7,13^ that PT is unobservable in the E148A mutant regardless of the presence of Cl^−^_cen_.

**Figure 6.**
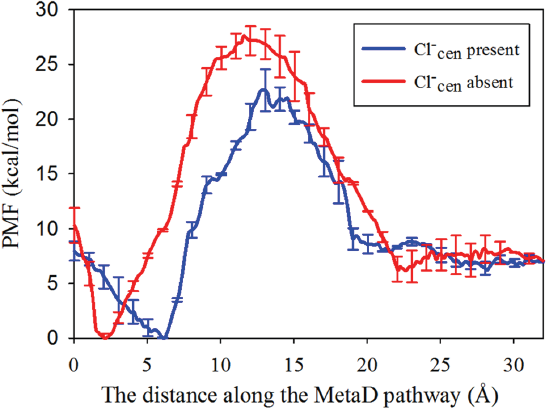
The PMF for PT in the E148A mutant, from E203 to the extracellular region, with Cl^−^_cen_ present (blue) and absent (red).

As previously noted, this mutant is especially intriguing due to the finding that H^+^ flux can be rescued by adding free glutamate to the solution in the absence of Cl^−^.^15^ Feng *et al.* solved the crystal structure for this mutant and found the carboxyl group of the glutamate from solution bound to the S_cen_ site. Its position was similar to the “down” conformation of the WT E148 with Cl^−^_cen_ absent shown in Figure 4C. We expect that the binding of the glutamate to S_cen_ in E148A mutant may be energetically less favorable than the down conformation of E148 in WT, due to steric hindrance between the substrate and the surrounding protein residues. Assuming that 1) the difference between two systems only locally affects the PMF for PT in E148A when the glutamate is bound to S_cen_ (corresponding to the E148 down confirmation in the WT PMF with Cl^−^_cen_ absent in Figure 3B), and 2) the binding of the glutamate in the E148A mutant is destabilized by ~3-4 kcal/mol compared to WT, decreasing the free energy barrier for PT via glutamate, then the rate constant would be ~150-780-fold greater than that in WT, allowing the H^+^ flux in the E148A mutant to be observed in experiment. As the glutamate ion binds less strongly to S_cen_ than Cl^−^, the free glutamate could only occupy this site in the absence of Cl^−^. This would explain the only observed PT through ClC-ec1 (in the E148A mutant + glutamate) in the absence of Cl^−^.

## Conclusions

Our multiscale simulations were performed to investigate the ClC-ec1 PT mechanism from E148 to the extracellular solution, with and without Cl^−^ bound at S_cen_. It was found to consist of two elementary steps: rotation of E148 from the “down” to “up” conformations, followed by deprotonation of E148 to the extracellular region. The two-step process was described by the curvilinear pathway followed by the excess proton, providing a single continuous CV that was sampled to collect a continuous PMF for this process.

Our calculations of the PT PMFs and the rate constants with either Cl^−^_cen_ present or absent suggest that a (perhaps the) key mechanism of Cl^−^/H^+^ coupling in ClC-ec1 is that Cl^−^_cen_ significantly facilitates the deprotonation of E148. For the outward flux with Cl^−^_cen_ present, the calculated effective rate constant for this two-step process was comparable to the experimentally observed overall PT rate, suggesting that this PT step is rate-limiting. When Cl^−^ _cen_ is absent, E148 is stabilized in the down conformation, bound to the S_cen_ site where further PT steps are effectively blocked and the calculated PT rate constant is below the experimentally measurable range.

The Cl^−^/H^+^ exchange mechanism can also operate in the opposite direction. For the inward H^+^ flux (the outward Cl^−^ flux), the rate-limiting step for the overall PT is likely PT from E148 to E203, which is also facilitated by Cl^−^_cen_. Thus, an essential molecular mechanism of the Cl^−^/H^+^ coupling is E148 rotation/deprotonation, which is facilitated by the presence of Cl^−^_cen_. In addition, the simulation structures at the up and the down conformations of E148 are consistent with several X-ray crystal structures showing the conformational change of E148. Furthermore, the pKa of E148 calculated from the PMF agrees well with the experimentally determined value.

It has been proposed that PT in ClC-ec1 could be coupled with other protein conformational changes, larger than the rotation of E148, outside of the central region. The crystal structures of ClC proteins have not revealed any large-scale conformational change among different structures, unlike other transporters.^1^ However, experimental^11,39,40,41^ and computational^41,42^ studies indicate conformational changes that are coupled with transport of Cl^−^ and H^+^, although the details of the changes are still uncertain. Although the results presented herein are not in conflict with these studies, they do suggest that one aspect of H^+^/Cl^−^ coupling (the dependence of PT on Cl^−^ occupancy) does not require larger conformational changes. Future studies that are able to provide information about the magnitude of the protein conformational change and its influence on ion flux, will further improve our understanding of this intriguing protein.

The PMF for PT was also calculated in E148A mutant, from E203 through the central region and to the extracellular solution. The free energy barrier for PT is increased compared to the WT protein when the proton passes through the narrow, dehydrated region around A148. The resulting PMFs showed that the free energy barrier for PT is high enough to reduce the PT in the E148A mutant to below detectable limits in both directions of the H^+^ flux, regardless of the presence of Cl^−^_cen_. The simulation results agree with the experimental findings for E148A mutant, where PT is not observed, although Cl^−^ can passively transit through the protein.

Collectively, our results suggest that the rate-limiting step for PT through ClC-ec1 requires the presence of Cl^−^_cen_ and depends on the direction of flow: for outward flux, the smallest calculated rate constant corresponds to E148 deprotonation to the extracellular solution, while for inward flux, the smallest rate constant comes from deprotonation of E148 to E203 in the central region. This work and previous studies have elucidated many elementary steps in the Cl^−^ /H^+^ exchange mechanism.^18,43^ Our future efforts will aim to determine how they combine to produce the macroscopically observable protein activity, such as the stoichiometric exchange ratio that remains consistent at different external ion concentrations.

## Supporting information

Additional details such as for the system setup and the procedure of the parameterization of MS-RMD model for E148, four figures, and one table are included in the supporting information. This material is available free of charge via the internet at http://pubs.acs.org.

## Acknowledgements

We thank Professor Christopher Miller of Brandeis University and Professor Alessio Accardi of Cornell University for their invaluable input on this work. The personnel in this research were supported by the National Institutes of Health (NIH Grant R01-GM053148). The computational resources in this research were provided by: the Extreme Science and Engineering Discovery Environment (XSEDE), which is supported by National Science Foundation grant number ACI-1053575; the U.S. Department of Defense (DOD) High Performance Computing Modernization Program at the Engineer Research and Development Center (ERDC) and Navy DOD Supercomputing Resource Centers; the University of Chicago Research Computing Center (RCC); and the NIH through resources provided by the Computation Institute and the Biological Sciences Division of the University of Chicago and Argonne National Laboratory, under grant 1S10OD018495-01.

---

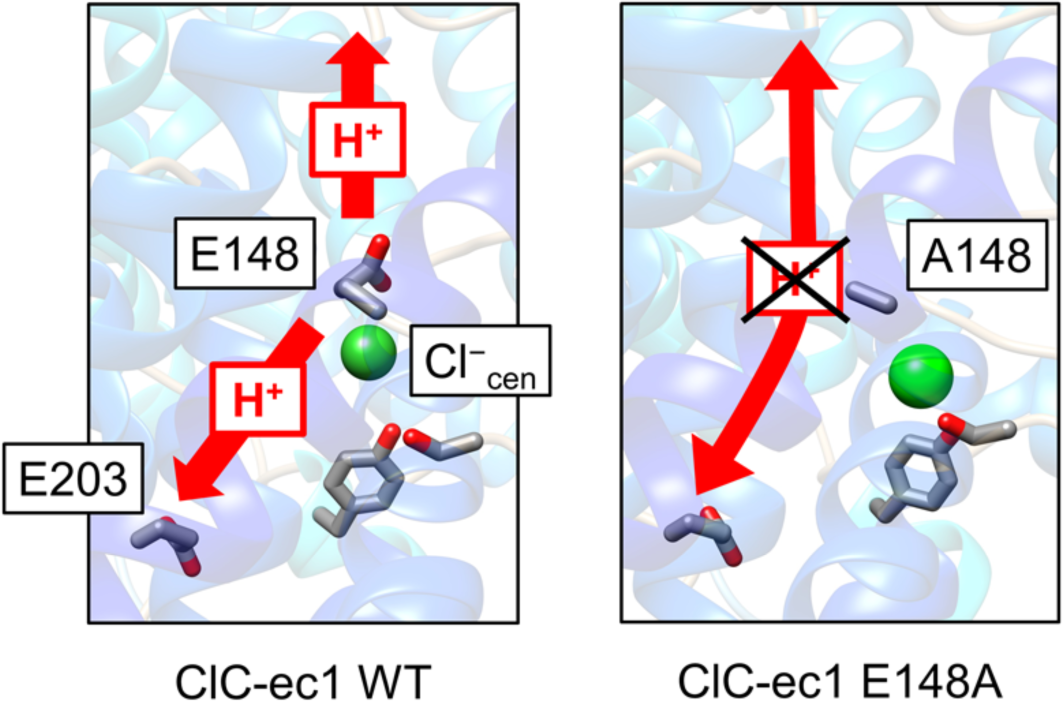

